# Characterizing cellular subpopulations critical to treatment response in autoimmune diseases

**DOI:** 10.64898/2026.01.05.696211

**Authors:** Siqi Sun, Mulini Pingili, Shweta Yadav, Zhuoya Wan, Michael Macoritto, Kathleen Smith, Jing Wang, Dan Chang, Naim A Mahi

## Abstract

Single-cell RNA sequencing provides a powerful approach for characterizing cell types, states, and lineages within heterogeneous tissues. However, identifying cell subpopulations that drive phenotypes, particularly treatment responses, remains a challenge. In this study, we performed comprehensive analyses to identify treatment response–associated cell subpopulations in autoimmune diseases by mapping bulk response information onto single-cell data. We integrated single-cell and bulk biopsy data from 314 responders and 619 non-responders treated with six therapeutics targeting tumor necrosis factor (TNF), integrin, or interleukin pathways in inflammatory bowel diseases (IBD) and psoriasis (PsO). Our analyses captured 128,428 interactions among 3,617 differentially expressed genes (DEGs), 852 pathways, and nine cell types spanning immune, stromal, and epithelial compartments. The importance of epithelial barrier integrity and enterocyte-mediated permeability in responses and inflammatory signaling in macrophages in non-responses in all tested therapies were highlighted by the presence of shared DEGs and pathways in both Crohn’s disease (CD) and ulcerative colitis (UC). In PsO, keratinocytes drove non-response to integrin-targeting therapies via disrupted adhesion, migration, and sustained epidermal inflammation. Additionally we introduce SCTRAD (https://immbioinfoabbv.shinyapps.io/SCTRAD/), an online web-based platform that allows exploration of mechanisms that influence the heterogeneity of cellular responses through DEGs and pathways at the single-cell level, as well as analysis of the cell-type-specific drug-related gene network. Our findings provide a comprehensive analysis of the role of cellular heterogeneity in treatment outcomes and for advancing precision medicine strategies in autoimmune diseases.

## 1. Introduction

Single-cell RNA sequencing (scRNA-seq) has emerged as a powerful tool to characterize cells at unprecedented resolution and uncover cellular mechanisms underlying complex diseases ^1,2^. This technology has the ability to reveal substantial cellular heterogeneity and demonstrate how aberrant cellular states can drive disease phenotypes ^3–7^. For example, in ulcerative colitis (UC), remodeling of the colonic cellular landscape has been observed, with enrichments of mast cells ^4^, CD8⁺IL-17⁺ T cells ^5^, and regulatory T (Treg) cells ^3,6^. Moreover, significant differences in immune cell and fibroblast compositions have been reported between infliximab responders and non-responders in inflammatory bowel disease (IBD) ^7^. Together, these findings highlight the critical role of cellular heterogeneity in shaping both disease progression and therapeutic outcomes.

Inflammatory bowel disease (IBD), primarily represented by Crohn’s disease (CD) and ulcerative colitis (UC), is characterized by chronic, relapsing intestinal inflammation ^8^. Currently in IBD, biologic therapies have advanced beyond symptomatic control, aiming instead to modify the disease course ^9,10^. These include anti–tumor necrosis factor (anti-TNF) agents such as infliximab and golimumab, which neutralize TNF-α, a central proinflammatory cytokine ^11^; anti-integrin agents such as vedolizumab and etrolizumab, which block lymphocyte trafficking by targeting α4β7 or αEβ7 integrins ^12,13^; and ustekinumab, which targets the shared p40 subunit of IL-12 and IL-23 ^14,15^. In addition, psoriasis (PsO), a chronic inflammatory and autoimmune skin disorder affecting over 60 million individuals globally ^16^. Psoriasis is characterized by erythematous, scaly plaques, potentially with pruritus, fissuring, and bleeding ^16^. Therapeutic management includes biologic agents such as ustekinumab and secukinumab, an anti–IL-17A monoclonal antibody ^14,17^. Despite their efficacy, cellular mechanisms underlying differential treatment outcomes including primary non-response and loss of response remain poorly understood.

Identifying specific cell subpopulations that drive clinical phenotypes such as treatment response is of fundamental importance, as it enables development of cell population–specific targeted therapies and facilitates discovery of clinically relevant biomarkers ^18,19^. However, identifying these subpopulations from single-cell data remains challenging. Generation of scRNA-seq data across large treatment cohorts is limited by cost and sample availability ^20^. Integrating single-cell data with additional phenotype-level information from bulk omics data provides a computational solution to identify biologically and clinically significant cell subpopulations ^20–22^. Such integrative methods have identified survival-associated cell populations in lung cancer and immunotherapy-responsive T cell states in melanoma ^20–22^. However, a large-scale, systematic characterization of the cellular subpopulations that influence treatment responses in autoimmune diseases, together with an integrated analysis of drug–gene interactions and the role of cellular heterogeneity in modulating therapeutic outcomes, is still needed^23^.

In this study, we integrated single-cell and bulk transcriptomic profiles with clinical response data from 933 patients (314 responders and 619 non-responders) treated with six biologic agents. Computational mapping of treatment response to single-cell populations identified cell subpopulations driving response and non-response to biologic therapies. Our analysis uncovered gene–pathway–cell relationships across diseases and treatments and highlighted both shared and distinct mechanisms of cellular heterogeneity on treatment response. The consolidated results are available for exploration in a Shiny-based web application “SCTRAD” (cell **S**ubpopulations **C**ritical to **T**reatment **R**esponse in **A**utoimmune **D**iseases) (https://immbioinfoabbv.shinyapps.io/SCTRAD/). These findings enable discovery of therapeutic targets and further advance precision medicine in immune-mediated diseases.

## 2. Materials and Methods

### 2.1 Dataset collection and pre-processing

We performed a comprehensive search of Gene Expression Omnibus (GEO) ^24^ and ArrayExpress ^25^ repositories for available CD, UC and PsO bulk transcriptomic studies deposited prior to April 2025 and selected 10 datasets with clinical response information to therapeutics (Infliximab ^26–30^, Golimumab ^31^, Secukinumab ^32^, Vedolizumab ^30^, Etrolizumab ^33,34^, Ustekinumab ^35–38^). The response criteria and study populations were defined as described in the original publications. Gene expression data were generated using both microarray and bulk RNA-Seq (Table. 1). Microarray data were RMA normalized, while RNA-seq data were normalized using the TMM method ^39,40^. Gene-level expression for each sample was calculated by averaging expression values from probes mapping to the same genes while excluding individual probes that were associated with more than one transcript, as previously described ^7,41^.

We curated autoimmune scRNA-seq datasets comprising patient biopsies from UC, CD, and PsO, all collected prior to biologic treatment. Additionally, we included a scRNA-seq dataset from adalimumab-treated IBD patients (TAURUS study). Several QC checks on the scRNA-seq were further performed, including removal of cells with abnormal mitochondrial (mt) RNA and gene levels (percent mt < 25 & nFeature_RNA > 200 & nFeature_RNA < 5000) ^7,41^. Cellular populations composed of at least 500 cells were retained and cells with fewer than 100 detected genes were excluded. Genes corresponding to HLA, immunoglobulin, RNA, mitochondrial (MT), and ribosomal protein (RP) categories (based on HUGO gene nomenclature) were removed. UMI counts were normalized by dividing the raw counts by total detected RNAs in each cell and multiplying by 10,000 (UMIs per million), followed by log-normalization ^7,41^. Highly variable genes were identified by selecting the top 3,000 genes based on mean expression and dispersion thresholds ^42,43^. Dimensionality reduction was performed using PCA on scaled data. A neighborhood graph was constructed using 30 nearest neighbors and 40 principal components, followed by UMAP embedding for visualization of cell clusters ^42,43^.

### 2.2 Identification of treatment-related cell types

The scRNA-seq datasets of untreated CD, UC and PsO patient intestine and skin biopsies respectively, were further annotated for cell types by GPTCelltype^44^ and evaluated by subject matter experts. Cell subpopulations associated with response and non-response were identified by integrating bulk and scRNA-seq data using Scissor ^20^. Scissor calculates a correlation matrix and a cell-cell similarity network using bulk RNA-seq profiles with known phenotypes and individual scRNA-seq cells. It then integrates these into a network-regularized sparse regression model to identify cell subpopulations positively or negatively associated with the phenotypes ^20^. By default, Scissor restricted the number of treatment-associated cells selected to no more than 20% of the total cells in the single-cell dataset ^20^. All other cells were considered as background or unselected cells. Cell types with at least 500 scissor-selected cells were included for the downstream analyses. They were classified as response-related if the number of responder cells were more than 1.5-fold higher than non-responders, and as non–response–related if non-responders were more than 1.5-fold higher than responders. For each cell type, percentages of response- and nonresponse-related cells were calculated relative to all treatment-related cells.

### 2.3 Gene and pathway analyses

For each cell type, treatment-related cells were compared to all other cells to identify differentially expressed genes (DEGs) using the findMarkers function from the Seurat package ^42,43^. DEGs with an adjusted p-value < 0.1 by the Wilcoxon rank-sum test were considered statistically significant ^45^. Pathway analysis on significant DEGs from cell types with at least 30 DEGs was performed using QIAGEN IPA ^46^. Significant canonical pathways were defined as those with a Benjamini–Hochberg adjusted p-value corresponding to –log₁₀(p) ≥ 1.3 and containing at least 5 DEGs.

### 2.4 Data availability

For bulk transcriptomic datasets, raw expression matrices were retrieved either from the original publications or GEO (Table 1). CD and UC scRNA-seq data can be downloaded from the single cell portal of the Broad Institute (https://singlecell.broadinstitute.org/single_cell) ^3,47^. PsO scRNA-seq data is available from GEO under accession number GSE162183 ^48^. Taurus raw data were downloaded from https://zenodo.org/records/14007626 ^49^. The UC spatial transcriptomics data is available from the original publication ^50^. The web-based platform SCTRAD, is available at https://immbioinfoabbv.shinyapps.io/SCTRAD/.

**Table. 1.** Bulk datasets with clinical response information.

| Datasets | ExperimentType | Disease | Source | Treatment | AgeGroup | Platform |
| --- | --- | --- | --- | --- | --- | --- |
| GSE92415 | Microarray | UC | Colon | Golimumab | Adult | Affymetrix HT HG-U133+ PM Array Plate |
| GSE73661 | Microarray | UC | Colon | Infliximab | Adult | Affymetrix Human Gene 1.0 ST Array |
| GSE73661 | Microarray | UC | Colon | Vedolizumab | Adult | Affymetrix Human Gene 1.0 ST Array |
| GSE73661 | Microarray | UC | Colon | Vedolizumab | Adult | Affymetrix Human Gene 1.0 ST Array |
| GSE73661 | Microarray | UC | Colon | Vedolizumab | Adult | Affymetrix Human Gene 1.0 ST Array |
| GSE72819 | RNASeq | UC | Colon | Etrolizumab | Adult | Illumina HiSeq 2000 |
| GSE23597 | Microarray | UC | Colon | Infliximab | Adult | Affymetrix Human Genome U133 Plus 2.0 |
| GSE206285 | Microarray | UC | Colon | Ustekinumab | Adult | Affymetrix HT HG-U133+ PM Array Plate |
| GSE16879 | Microarray | UC | Colon | Infliximab | Adult | Affymetrix Human Genome U133 Plus 2.0 |
| GSE16879 | Microarray | CD | Colon | Infliximab | Adult | Affymetrix Human Genome U133 Plus 2.0 |
| GSE152316 | RNASeq | CD | Colon | Etrolizumab | Adult | Illumina HiSeq 4000 |
| GSE132195 | RNASeq | PSO | Skin | Secukinumab | Adult | Illumina HiSeq 3000 |
| GSE12251 | Microarray | UC | Colon | Infliximab | Adult | Affymetrix Human Genome U133 Plus 2.0 |
| GSE117239 | Microarray | PSO | Skin | Etanercept | Adult | Affymetrix Human Genome U133 Plus 2.0 Array |
| GSE117239 | Microarray | PSO | Skin | Ustekinumab | Adult | Affymetrix Human Genome U133 Plus 2.0 Array |

| Timepoints | Number | Number | PMID_Ref |
| --- | --- | --- | --- |
| W6 | 61 | 48 | PMID: 23735746 |
| W6 | 8 | 15 | PMID:27802155 |
| W6 | 3 | 21 | PMID:27802155 |
| W12 | 3 | 10 | PMID:27802155 |
| W52 | 12 | 5 | PMID:27802155 |
| W10 | 3 | 40 | PMID: 26522261 |
| W8 | 63 | 18 | PMID: 21448149, PMID: 31039157 |
| W8 | 49 | 315 | PMID: 36192482, PMID: 36170836, PMID: 36109152 |
| W4-6 | 16 | 32 | PMID: 19956723 |
| W4_6 | 23 | 14 | PMID: 19956723 |
| W14 | 6 | 65 | PMID: 34467254 |
| W16 | 1 | 4 | PMID: 31715177 |
| W8 | 12 | 11 | PMID: 19700435 |
| W12 | 20 | 10 | PMID: 30703387 |
| W12 | 34 | 11 | PMID: 30703387 |

## 3. Results

### 3.1 Cell subpopulations in response to anti-TNF therapies in UC

We first evaluated the impact of infliximab, a chimeric monoclonal anti-TNF antibody, on treatment-associated cellular heterogeneity using transcriptomic data from 48 UC subjects (GSE16879) with annotated therapeutic response (Fig. 1). We identified 4,457 response-associated cells and 15,526 non–response-associated cells among a total of 114,512 cells, representing 17.4% of all Scissor-selected cells relative to the total cell population (Fig. 2A, Table. S1). Response-associated cells were epithelial populations: colonocytes, enterocytes, and goblet cells, whereas non–response-associated cells were distributed across most subpopulations, including B cells, cytotoxic T cells, endothelial cells, fibroblasts, macrophages, mast cells, plasma cells, proliferating cells, and T cells (Fig. 2A, 2B). These findings were consistently replicated across separate, independent infliximab UC datasets (Fig. S1), suggesting a central role of enterocytes and macrophages in mediating response and non-response, respectively.

**Fig. 1.**
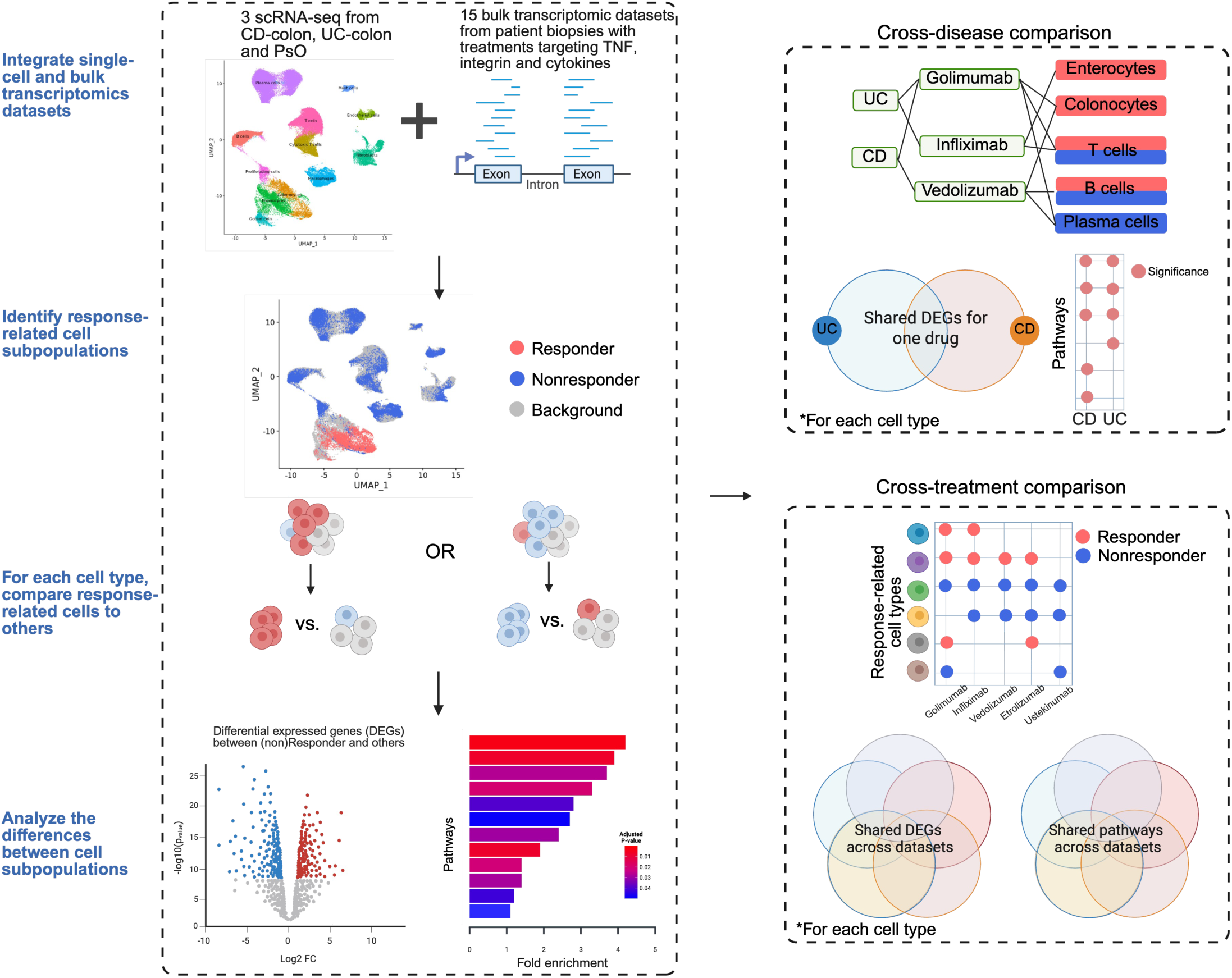
Analysis workflow. We processed scRNA-seq data from patients with IBD and PsO, along with bulk RNA-seq data from tissue biopsies treated with therapeutics targeting TNF, integrins, and cytokines. By integrating single-cell and bulk transcriptomic data, we identified treatment response–associated cell subpopulations for each condition. Within each cell type, dominant subpopulations (e.g., responders (R) or non-responders (NR)) were compared to other cells through differential gene expression and pathway analyses. Finally, we conducted cross-disease and cross-treatment comparisons to uncover shared and distinct molecular features.

**Fig. 2.**
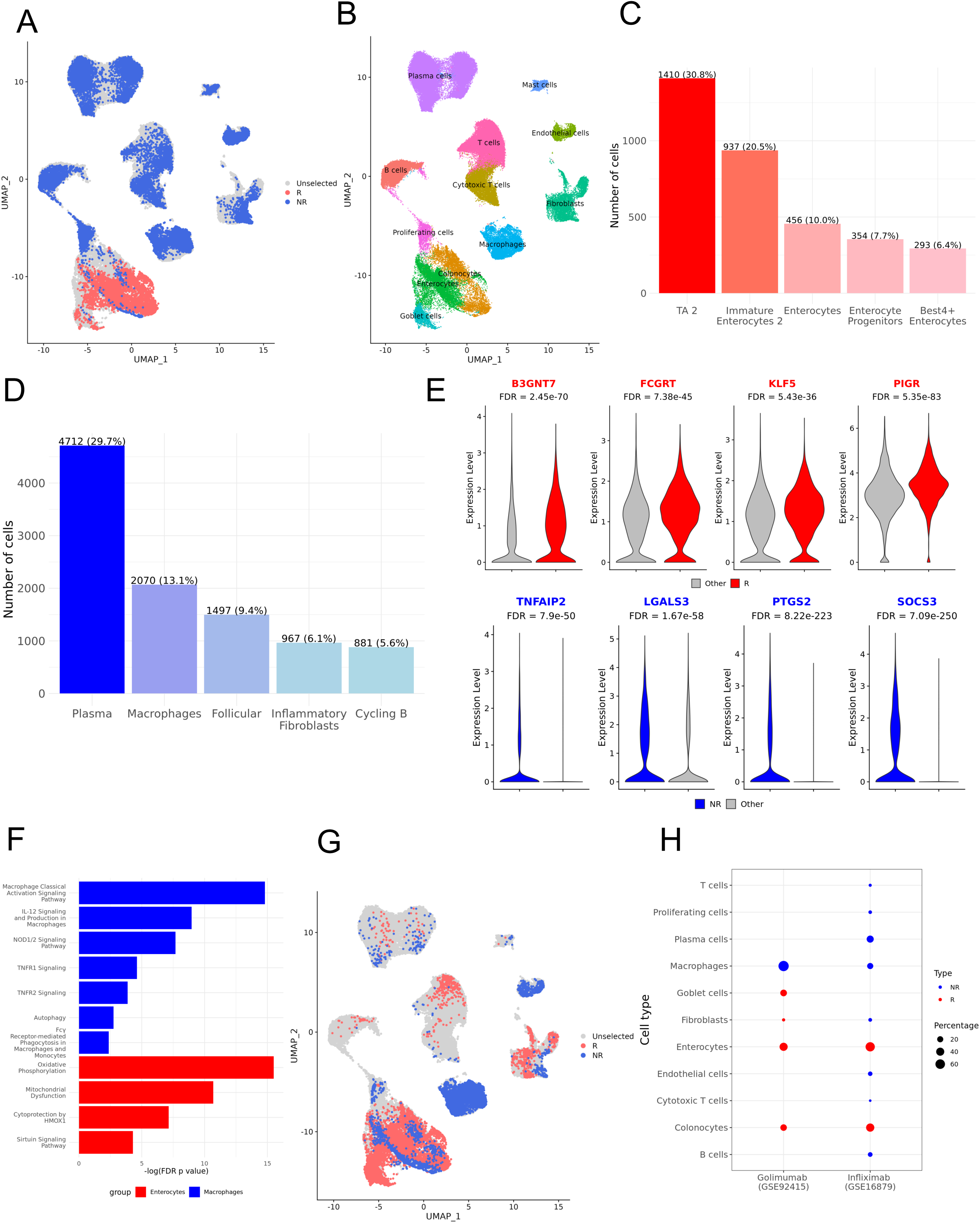
Cell subpopulations related to anti-TNF response in UC. (A) UMAP visualization of the response-related cells treated with infliximab in GSE16879. The red and blue dots are response and non-response-related cells, respectively. (B) The UMAP visualization of 114,512 UC cells. (C) The composition of response-related cells in the top five most abundant cell types. Values represent the number and the percentage among all response-related cells. (D) The composition of non-response-related cells in the top five most abundant cell types. Values represent the number and the percentage among all non-response-related cells. (E) The expression levels of key genes in response-related cells (red) and non-response-related cells (blue). The FDR was the adjusted P value calculated by the two-tailed Wilcoxon rank sum test. (F) The representative pathways significantly enriched in response-related (red) or non-response-related (blue) cells compared to all other cells (FDR <0.05). (G) UMAP visualization of the response-related cells treated with Golimumab in GSE92415. (H) The composition of different cell types relative to the total number of response-related (red) or non-response-related (blue) cells.

At higher resolution (cell types defined by the original publication), treatment response was specifically associated with distinct epithelial subpopulations, including 1410 absorptive transit amplifying (TA) cells 2 (30.8% of all response-related cells), followed by 937 immature enterocytes, and 456 enterocytes (Fig. 2C). In contrast, non-response was dominated by 4,712 plasma cells (29.7% of all nonresponse-related cells), 2,070 macrophages and 1,497 follicular cells (Fig. 2D). Transcriptional profiling of response enterocytes relative to all other enterocytes revealed 230 differentially expressed genes, such as *KLF5* (FDR=5.43e-36) and *PIGR* (FDR=5.35e-83) (Fig. 2E). Specifically, *KLF5* is a transcription factor highly expressed in intestinal epithelial cells and plays a critical role in preserving epithelial integrity and protecting against Th17-mediated immune and inflammatory responses ^51^. In addition, the polymeric immunoglobulin receptor (pIgR) mediates the transcytosis of IgA and IgM into the intestinal lumen to inhibit pathogen colonization for gut microbiota hemostasis ^52^. We observed significant enriched pathways, such as the sirtuin signaling pathway (FDR=5e-5) (Fig. 2F), which plays a protective role in maintaining gut homeostasis by reducing the expression of proinflammatory genes and modulating antimicrobial protein levels in the small intestine ^53–55^. In contrast, non-response-associated macrophages exhibited 457 differentially expressed genes, such as *TNFAIP2* (FDR=7.9e-50) and *SOCS3* (FDR=7.09e-250) (Fig. 2E). TNFα-induced protein 2 (TNFAIP2), upregulated under TNFα stimulation, has been implicated in promoting angiogenesis, enhancing cell proliferation, adhesion, and migration, as well as inducing tunneling nanotube formation during tumor invasion ^56^. In addition, the suppressors of cytokine signaling (SOCS) 3 protein is the key physiological regulators of cytokine-mediated STAT3 signaling and M2 differentiation in IBD^57^. Pathways such as TNFR1 signaling (FDR=2.4e-5) and classical macrophage activation pathways are significantly enriched (FDR=1.58e-15) (Fig. 2F).

When comparing two anti-TNF therapeutics, golimumab versus infliximab, enterocytes and macrophages continued to mediate response and non-response, respectively, but the most pronounced differences between the two treatments were observed in fibroblast populations (Fig. 2G, 2H). Fibroblasts were predominantly non–response-associated in infliximab-treated samples, comprising 1,180 cells (7.6% of all non–response-associated cells) in GSE16879 (Table S1). In contrast, in golimumab-treated samples, fibroblasts were mainly response-associated, comprising 482 cells (6.6% of all response-associated cells) (Fig. 2H, Table. S1). This may be due to distinct signaling pathways in fibroblasts that drive differential responses. For example, TNFR2 non-canonical NF-κB pathway (FDR= 3.98e-12) and TNFR1 signaling (FDR= 3.16e-02) were observed in response to infliximab but not to golimumab (Table. S2).

### 3.2 Cell subpopulation in response to anti-integrins in UC

Using the same single-cell dataset from subjects from UC, we compared cellular responses to two anti-integrin therapeutics: vedolizumab and etrolizumab. Following vedolizumab treatment, 11,403 cells were associated with response, and were predominantly colonocytes and enterocytes. In contrast, 8,764 cells were linked to non-response, with enrichment in endothelial cells, fibroblasts, and macrophages (Fig. 3A, 3C, Table. S1). For etrolizumab, 4,841 response-associated cells comprised colonocytes and fibroblasts, while 12,766 non-response-associated cells included B cells, T cells, cytotoxic T cells, and macrophages. (Fig. 3B, 3C, Table. S1). Notably, enterocytes and colonocytes consistently associated with response, whereas macrophages were consistently linked to non-response. Fibroblasts showed treatment-specific associations: they were largely non-response associated in vedolizumab (5,062 cells) but mostly response-associated in etrolizumab (3,924 cells) (Fig. 3A-C, Table. S1). Transcriptional profiling shows 439 shared DEGs in the response colonocytes between vedolizumab and etrolizumab, such as *PIGR* (Fig. 3D). Pathways such as tight junction signaling and cell junction organization are enriched significantly (Fig. 4D). In contrast, only 4 shared DEGs (*EREG*, *IL1RN*, *NFKBIA*, and *SRGN*) in the non-response macrophages between vedolizumab and etrolizumab (Fig. 3E). Specifically, *EREG* (epiregulin) promotes epithelial repair and regeneration, supporting mucosal healing ^58^. *IL1RN* encodes the interleukin-1 receptor antagonist (IL-1ra), which helps maintain immune homeostasis in the gut ^59,60^. *NFKBIA* encodes IκBα, an inhibitor of NF-κB, thereby modulating inflammatory gene expression and regulating mucosal inflammation^61^. *SRGN* (serglycin) is essential for the formation of mast cell secretory granules and mediates storage and secretion of proteases, chemokines, and cytokines, contributing to immune regulation ^62^. We also observed that 91 significantly enriched pathways are common, highlighting the importance of interleukin pathways (Fig. 4F).

**Fig. 3.**
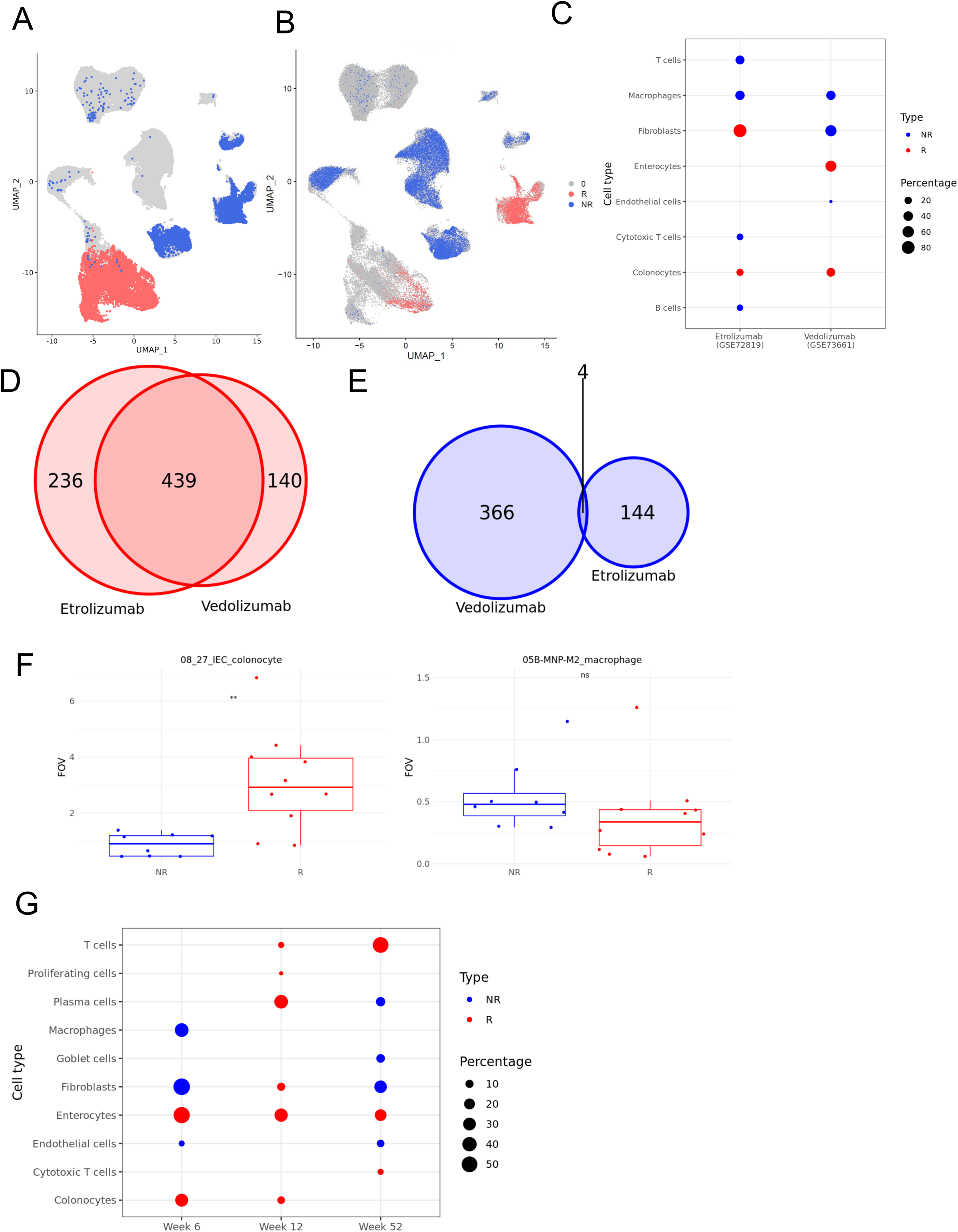
Cell subpopulations related to anti-integrin response in UC. (A) UMAP visualization of the response-related cells treated with Vedolizumab in GSE73661. (B) UMAP visualization of the response-related cells treated with etrolizumab in GSE72819. (C) The composition of different cell types relative to the total number of response-related (red) or non-response-related (blue) cells in response to different anti-integrins. (D) Common differentially expressed genes (DEGs) in R-dominant colonocytes across different anti-integrin treatments. (E) Common DEGs in NR-dominant macrophages across different anti-integrin treatments. (F) Comparison of cell abundance of colonocytes and macrophages in vedolizumab responders (R) and non-responders (NR) using independent spatial CosMx data. Data are presented as mean values in each FOV. (G)The composition of different cell types relative to the total number of response-related (red) or non-response-related (blue) cells in response to Vedolizumab at different timepoints.

**Fig. 4.**
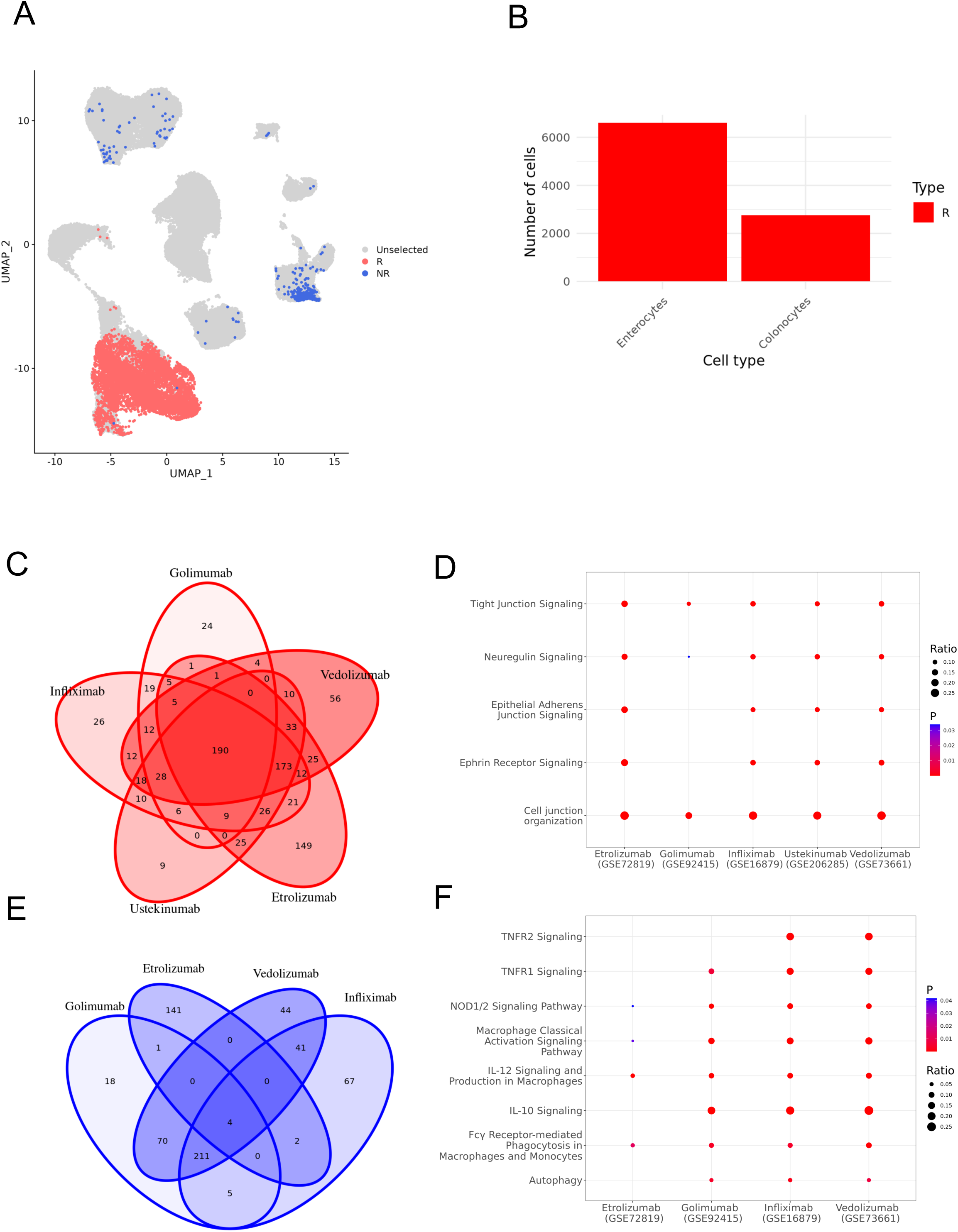
Cell subpopulations related to response to different therapeutics in UC. (A) UMAP visualization of the response-related cells treated with Vedolizumab in GSE73661. (B) Cell numbers of treatment-related cell types. (C) Common DEGs in R-dominant enterocytes across different biologic treatments. (D) Representative enriched pathways in R-dominant enterocytes across different biologic treatments. (E) Common DEGs in NR-dominant macrophages across different biologic treatments. (F) Representative enriched pathways in NR-dominant macrophages across different biologic treatments.

To further validate our findings regarding the response to vedolizumab, we incorporated two additional datasets to examine spatial localization and temporal dynamics at multiple timepoints during treatment, specifically within enterocytes and macrophages. The independent CosMx spatial dataset revealed significantly higher levels of colonocyte in responder patients (mean: 0.86 in NR vs. 3.12 in R, p < 0.01), whereas 05B-MNP-M2_macrophage abundance showed a trend toward being lower in non-responders (mean: 0.55 in NR vs. 0.38 in R) (Fig. 3F). In addition, several cell types exhibited dynamic and time-dependent associations with treatment outcome, potentially reflecting dose adjustments or progressive tissue remodeling following therapy ^3,30,63^. For example, among enterocyte subpopulations associated with response, we observed a gradual decline in cell numbers over time from 7,267 at week 6 to 6,310 at week 12 and 2,632 at week 52 (Fig. 3G, Table. S1). Fibroblasts were enriched in non-responders at week 6, shifted to responder-associated at week 12, and reverted to non-responder-associated by week 52 (Fig. 3G), highlighting their plasticity and context-dependent roles in inflammation and tissue repair ^3,63^.

### 3.3 Comparison of cell subpopulations in response to anti-TNFs, integrins and IL-12/23 in UC

In ustekinumab-treated patients, we identified 9,357 response-associated cells that were enriched in colonocytes and enterocytes with few non-response-associated cells identified (Fig. 4A, 4B).

Analyses of patients treated with diverse therapeutics, including anti-TNF agents (golimumab and infliximab), anti-integrins (vedolizumab and etrolizumab), and the anti–IL-12/23 antibody ustekinumab, revealed consistent patterns of response- and non–response-associated cell subpopulations despite targeting distinct pathways. Notably, enterocytes and colonocytes were recurrently enriched among response-associated cells, whereas macrophages were consistently enriched among non–response-associated cells (Fig. 2A, 3A, 3B, 4A, S2, Table. S1).

Further transcriptional analyses suggested response-associated colonocyte subpopulations shared 190 differentially expressed genes across treatment groups, including *IL23*, *CDH1*, and *ECM1* (Fig. 4C). Extracellular matrix protein-1 (ECM1) regulates colonic inflammation by controlling M1 macrophage polarization via the GM-CSF/STAT5 axis ^64^. In addition, it has shown that increased E-cadherin gene (*CDH1*) methylation in inflamed ileal mucosa links it to ileal inflammation and epithelial barrier dysfunction ^65,66^. Functional enrichment analysis of these genes identified 102 common pathways, such as tight junction signaling and cell junction organization, highlighting epithelial barrier restoration as a core mechanism of therapeutic response (Fig. 4D). Additionally, comparative analysis of non–response-related macrophage populations across all datasets revealed four consistently upregulated genes: *EREG*, *IL1RN*, *NFKBIA*, and *SRGN* (Fig. 4E). These genes support the significantly enriched inflammatory pathways, such as activin/inhibin signaling that drives inflammation and epithelial apoptosis in colitis ^67^, and the acute phase response, suggesting conserved transcriptional programs that underpin treatment resistance (Fig. 4F).

### 3.4 Comparison of cell subpopulations in response to anti-TNFs in CD and UC

Across 18 cell types in CD, we identified 2,256 infliximab response-associated cells and 1,562 non–response-associated cells, representing 11.9% of the total 32,022 cells (Fig. 5A, Table. S1). Response-associated cells were predominantly enterocytes, goblet cells, and Paneth cells, whereas non–response-associated cells were enriched in macrophages and fibroblasts (Fig. 5B, 5C). At a finer resolution, response was associated with 434 CA1⁺CA2⁺CA4⁻ enterocytes, 583 TMIGD1⁺MEP1A⁺ enterocytes, and 183 cycling epithelial cells (Table. S3). In contrast, non-response was primarily driven by 643 S100A8⁺S100A9⁺ monocytes and LYVE1+ macrophages (Table. S3). The cell subpopulations associated with infliximab response were consistent with those observed in UC (Fig. 2A–2D).

**Fig. 5.**
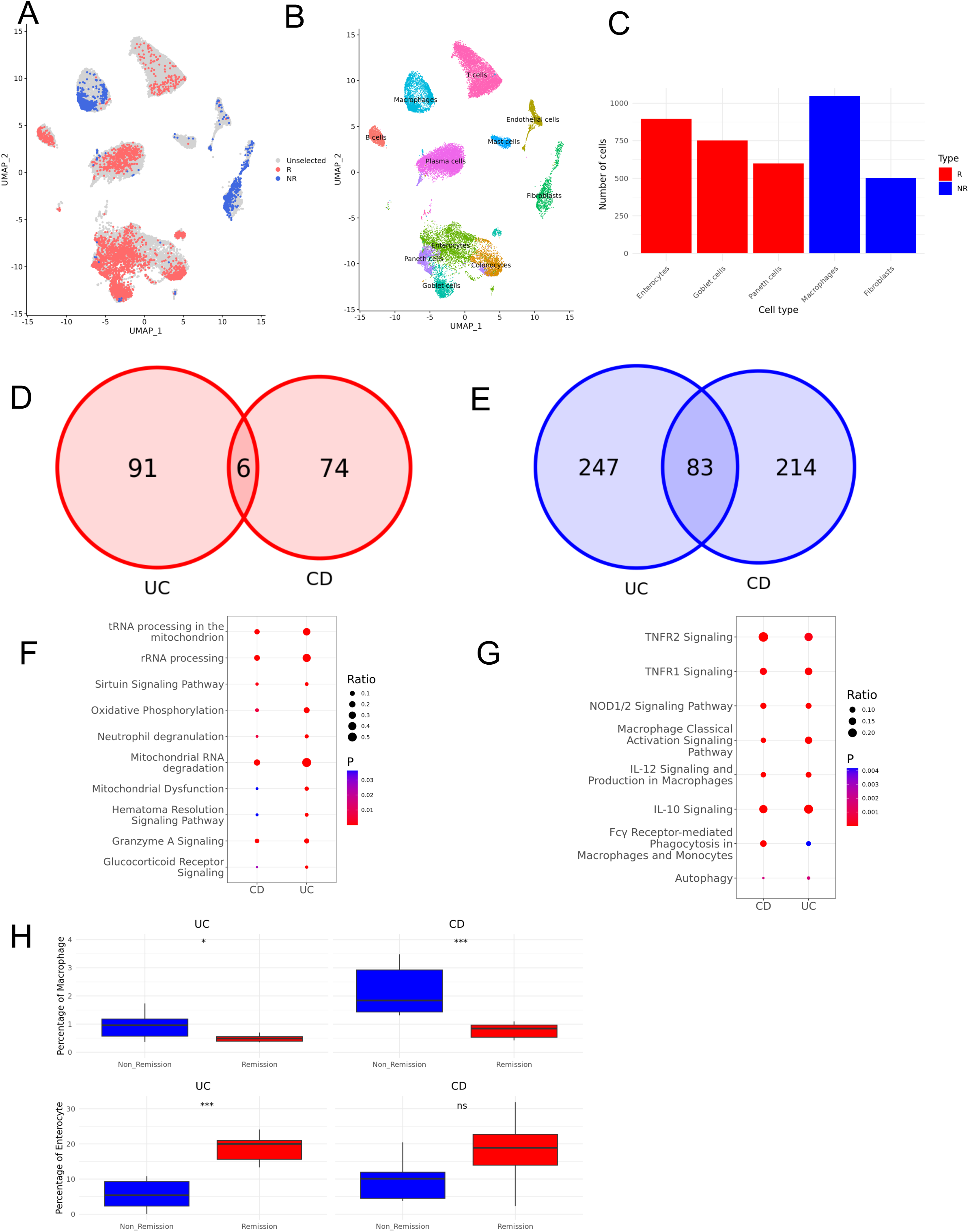
Cell subpopulations related to infliximab response in CD and UC. (A) UMAP visualization of the response-related cells treated with infliximab in GSE16879 in CD. (B) The UMAP visualization of 32,022 CD cells. (C) Cell numbers of treatment-related cell types in CD. (D) Common DEGs in R-dominant enterocytes between CD and UC. (E) Common DEGs in NR-dominant macrophages between CD and UC. (F) Representative enriched common pathways in R-dominant enterocytes between CD and UC. (G) Representative common enriched pathways in NR-dominant macrophages between CD and UC. (H) Cellular proportions of macrophages and enterocytes among all cells in remission and non-remission adalimumab-treated patients with CD and UC. Significance: ***p < 0.001, \**\**p < 0.01, *p < 0.05, ns > 0.05.

Across CD and UC, enterocytes from responders shared 10 pathways and 6 DEGs (Fig. 5D, 5F). Notably, common DEGs included *B3GNT7*, a glycosyltransferase essential for mucin biosynthesis and the maintenance of intestinal homeostasis ^68^. Common enriched pathways included the sirtuin signaling pathway, mitochondrial dysfunction and oxidative phosphorylation (Fig. 5F). It has been shown that mitochondrial mass and the expression of mitochondrial oxidative phosphorylation (OXPHOS) complexes in intestinal crypts tend to be reduced in IBD patients ^69^. By contrast, macrophages from non-responders shared 223 pathways and 83 DEGs, including *FCGR2A* and *IL8*, linked to TNFR2 signaling and classical macrophage activation (Fig. 5E, 5G). For example, *FCGR2A* mediates innate and cell-mediated immunity, promoting abundant secretion of stimulatory cytokines that drive T-regulatory cell–induced autophagy and antimicrobial activity ^70,71^, while *IL8* recruits neutrophils to the intestine, leading to iNOS and MMP release that contributes to epithelial cell damage ^72^.

Further analyses of single-cell data from the TAURUS study (adalimumab) revealed significant differences in cellular composition between remission and non-remission samples in IBD. In UC, the number of enterocytes were increased in remission compared to non-remission (18.8% vs. 7.94%, FDR<0.001), whereas macrophages were 0.49% in remission versus 1.12% in non-remission (FDR<0.05) (Fig. 5H). Similar trends were observed in CD patients (Fig. 5H). This further highlighted common disease mechanisms and therapeutic responses to anti-TNFs irrespective of disease subtype.

### 3.5 Cell subpopulations in responses to different anti-cytokine therapeutics in PsO

To further characterize the cellular subpopulations associated with therapeutic responses, we examined them in psoriasis (PsO). Across 12 cell types in psoriasis, the analysis of Ustekinumab-treated patients revealed 711 response-associated cells and 1,415 non–response-associated cells among 17,684 total cells (Fig. 6A, 6C, Table. S1). Response-associated cells were predominantly dermal fibroblasts, whereas non–response-associated cells were enriched in keratinocytes (Fig. 6D). In comparison, Secukinumab-treated patients exhibited 2,338 non–response-associated cells primarily enriched in basal keratinocytes, and keratinocytes (Fig. 6B-6D). Keratinocyte subpopulations consistently dominated the non-response landscape across both treatments, underscoring their key role in psoriasis persistence and treatment resistance (Fig. 6A-6D). Molecular analyses revealed 4 shared DEGs, including *KRT14, KRT5, MX1*, and *PTTG1*, which are involved keratinocyte structure, and differentiation, with an impact on cell mechanics, homeostasis and epidermal barrier function ^73–75^ (Fig. 6E). In addition, common pathways such as RhoA and MAPK signaling, which regulates cytoskeletal organization, cell migration, and inflammatory signaling in keratinocytes ^76,77^, were enriched, suggesting convergent mechanisms of treatment resistance across both therapeutics (Fig. 6F).

**Fig. 6.**
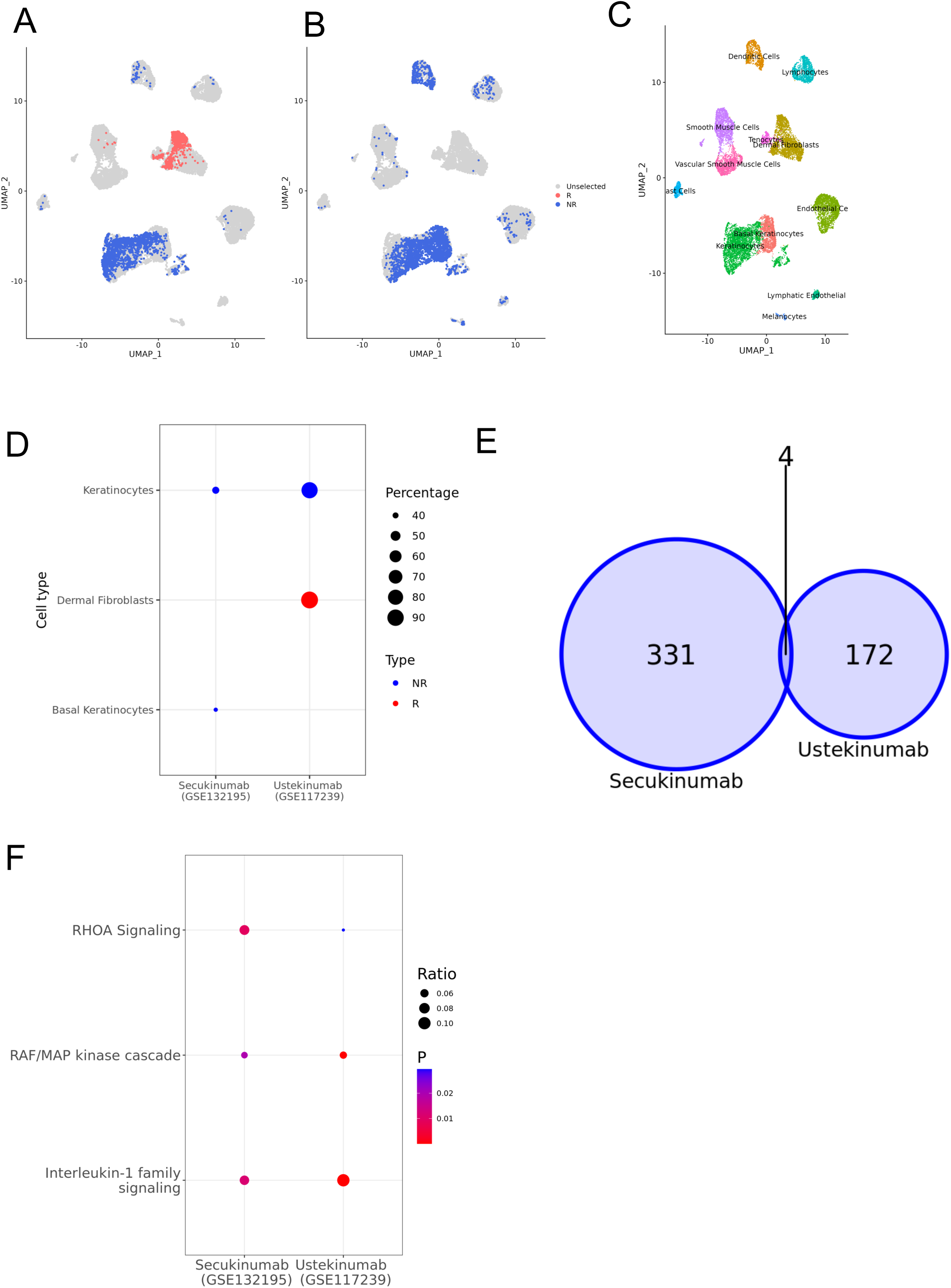
Cell subpopulations related to anti-integrin response in PsO. (A) UMAP visualization of the response-related cells treated with ustekinumab in GSE117239. (B) UMAP visualization of the response-related cells treated with secukinumab in GSE132195. (C) The UMAP visualization of 17,546 PsO cells. (D) The scatter plot shows the composition of different cell types relative to the total number of response-related (red) or non-response-related (blue) cells in response to different anti-integrins. (E) Common DEGs in NR-dominant keratinocytes across different anti-interleukin treatments. (F) Representative common enriched pathways in NR-dominant keratinocytes across different anti-integrin treatments.

## 4. Discussion

Identifying phenotype-specific cell subpopulations from single-cell data can provide critical insights into molecular programs and cellular heterogeneity that shape disease phenotypes and drug responses. However, a comprehensive characterization of the treatment-related cell subpopulations driving therapeutic responses in autoimmune diseases remains lacking. In this study, we mapped 933 bulk clinical samples with responses onto single-cell datasets and identified cellular signatures underlying treatment outcomes in IBD and PsO. Across different therapeutic classes (TNF inhibitors, other cytokine blockers, or integrin-targeting agents) and IBD subtypes (CD and UC), enterocyte subpopulations were consistently associated with positive treatment response, whereas macrophage subpopulations were consistently linked to non-response. Furthermore, the integration of independent single-cell transcriptomics data revealed the temporal dynamics and spatial distribution of those critical cell subpopulations, and collectively validated the results. The shared DEGs and pathways underscore the critical role of epithelial barrier integrity and mucosal healing in mediating treatment response, as well as the contribution of inflammatory signaling and cytokine/chemokine production in macrophages to non-response. Similarly, recent studies in IBD have identified that one patient subtype “S2” exhibits higher response rates to biologic therapies (corticosteroids, infliximab, vedolizumab, and ustekinumab) with enrichment of epithelial cells including immature enterocytes, compared with subtype “S1” which is enriched in immune and stromal cells such as inflammatory monocytes and inflammatory fibroblasts ^78^. Sun et al. also identified the presence of significantly more absorptive cells, including distinct enterocyte subtypes, and fewer monocytes in responders to infliximab compared to non-responders ^7^.

Additionally, in PsO, keratinocytes were associated with non-response to integrin-targeting agents, suggesting that aberrant keratinocyte proliferation and differentiation may impair therapeutic efficacy through the epidermis (Fig. 6) This is in line with prior studies highlighting excessive growth and aberrant differentiation of keratinocytes as a hallmark of psoriasis ^79^. This shared resistance across both therapeutics highlights the complex interplay between keratinocytes and immune cells, particularly through inflammatory processes regulated by signaling pathways such as the RAF/MAP kinase cascade and interleukin-1 family signaling, as previously reported (Fig. 6) ^80,81^.

Fibroblasts exhibit remarkable plasticity, displaying context-dependent roles in response to therapeutics targeting IBD (Fig. 2-4). Their effects vary even among agents targeting similar pathways (Fig. 2, 3). For example, fibroblasts contribute to the therapeutic response to golimumab but are linked to nonresponse with infliximab, despite both targeting TNF-α (Fig. 2). This complements previous studies in infliximab: Smillie et al^3^ reported that the infliximab resistance signature was strongly enriched in inflammatory fibroblasts in UC, and Martin et al. ^82^ reached a similar conclusion in CD. Similarly, fibroblasts are associated with a favorable response to etrolizumab (anti-β7 integrin) whereas responses to vedolizumab (anti-α4β7 integrin) are limited (Fig. 3). These contrasting effects may be explained by the heterogeneity of fibroblast populations, particularly inflammation-associated fibroblasts, which execute distinct inflammatory functions and orchestrate complex cell–cell interactions ^83^.

One limitation of this study is the paucity of single-cell and bulk RNA-seq data, which constrains the breadth of drug classes and disease contexts that can be analyzed. In the future, as more transcriptomic datasets become available across diverse therapeutic areas, we expect to identify and characterize a broader range of treatment-critical cell types, along with a more precise quantification of their association strengths. In addition, the integration of other modalities of single-cell data, such as scATAC-seq and single-cell proteomics, would further reveal regulatory networks or signaling interactions within those critical cell types, thereby providing additional insights into disease mechanisms and treatment responses ^84^.

In this study, we have identified cell subpopulations that are critical for responses to therapeutics targeting TNFand other cytokine pathways and integrins in multiple autoimmune diseases and explored their underlying cellular mechanisms. Enterocytes were consistently associated with therapeutic response, whereas macrophages were linked to non-response, independent of disease subtype (UC or CD) or biologic agent. In contrast, fibroblasts and T cells exhibited treatment- and time-dependent responses. Additionally, in psoriasis, keratinocytes were identified as drivers of non-response to integrin-targeting therapies. All analyses have been deposited in SCTRAD (https://immbioinfoabbv.shinyapps.io/SCTRAD/), an interactive platform that allows users to explore relationships among therapeutics, genes, pathways, cellular responses, cell types and diseases. Our findings highlight the potential of leveraging scRNA-seq for drug repurposing by pinpointing specific cell subpopulations central to treatment responses, thereby advancing precision medicine in autoimmune diseases.

## Funding

Financial support for this research was provided by AbbVie.

## Acknowledgements

We would like to acknowledge the valuable feedback from Archana Iyer, Charles Lu, Sarah Kongpachith, and Fedik Rahimov.

---

**Fig. S1** Cell subpopulations related to infliximab responses in UC across datasets.

**Fig. S2** Cell subpopulations related to different biologic responses in UC across datasets.

## Competing interests

The authors declare no competing interests.

## Disclosure statement

All Authors were or are employees of AbbVie at the time of the study. The design, study conduct, and financial support for this research were provided by AbbVie. AbbVie participated in the interpretation of data, review, and approval of the publication. No honoraria or payments were made for authorship.

## Authors’ contributions

Conceptualization: JW, DC, SS; Data curation: SS, SY, MP, ZW; Formal analysis: SS, SY, MP, ZW; Project Administration: JW, DC, NM; Writing: SS, ZW, MM, KS, NM.

## Notes

https://immbioinfoabbv.shinyapps.io/SCTRAD/

### Competing Interest Statement

The authors have declared no competing interest.

