## Supplementary figures and images for "Characterizing cellular subpopulations critical to treatment response in autoimmune diseases"

### Fig. S1-Fig. S2

#### Slide 1
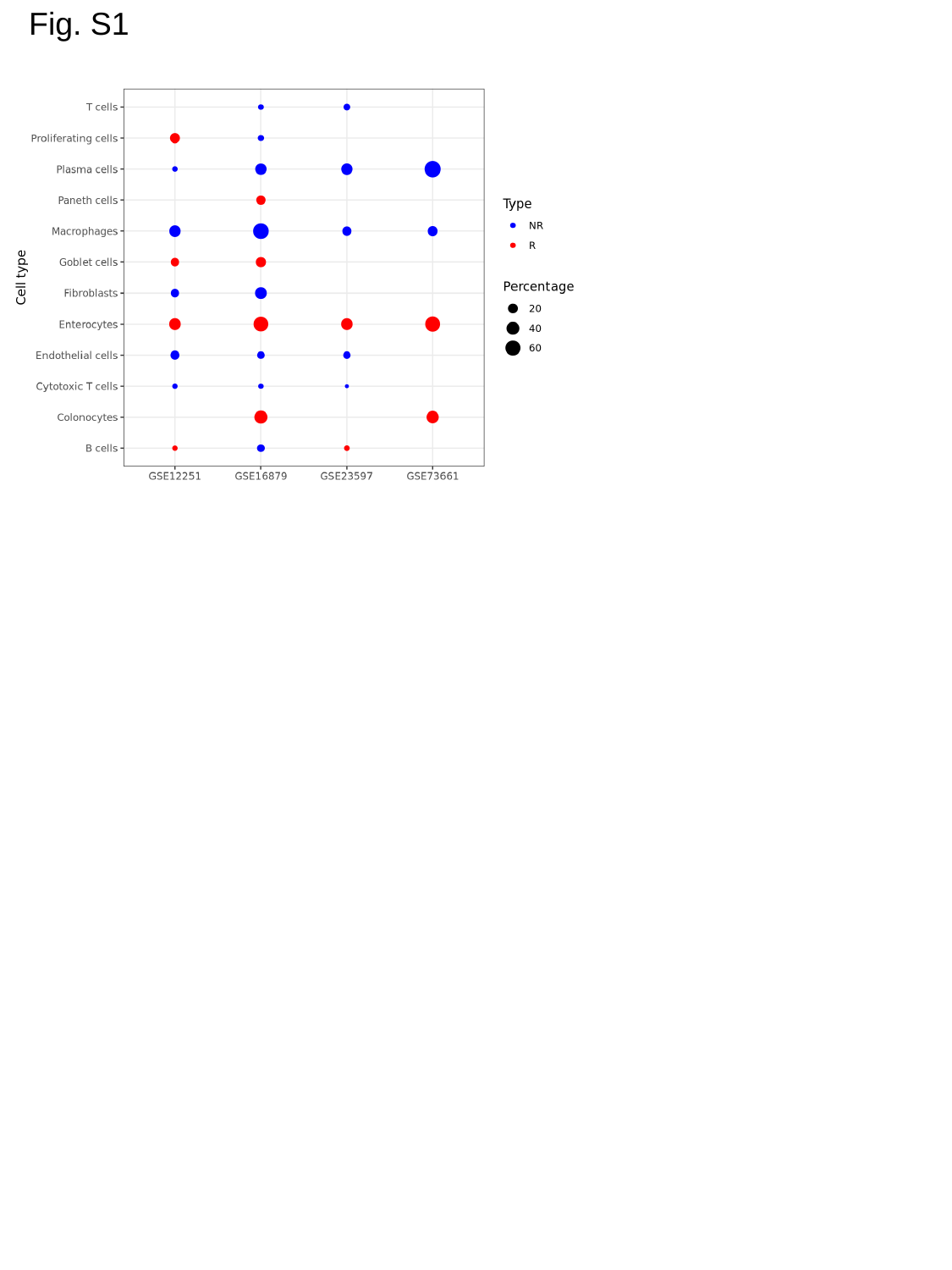

Fig. S1

#### Slide 2
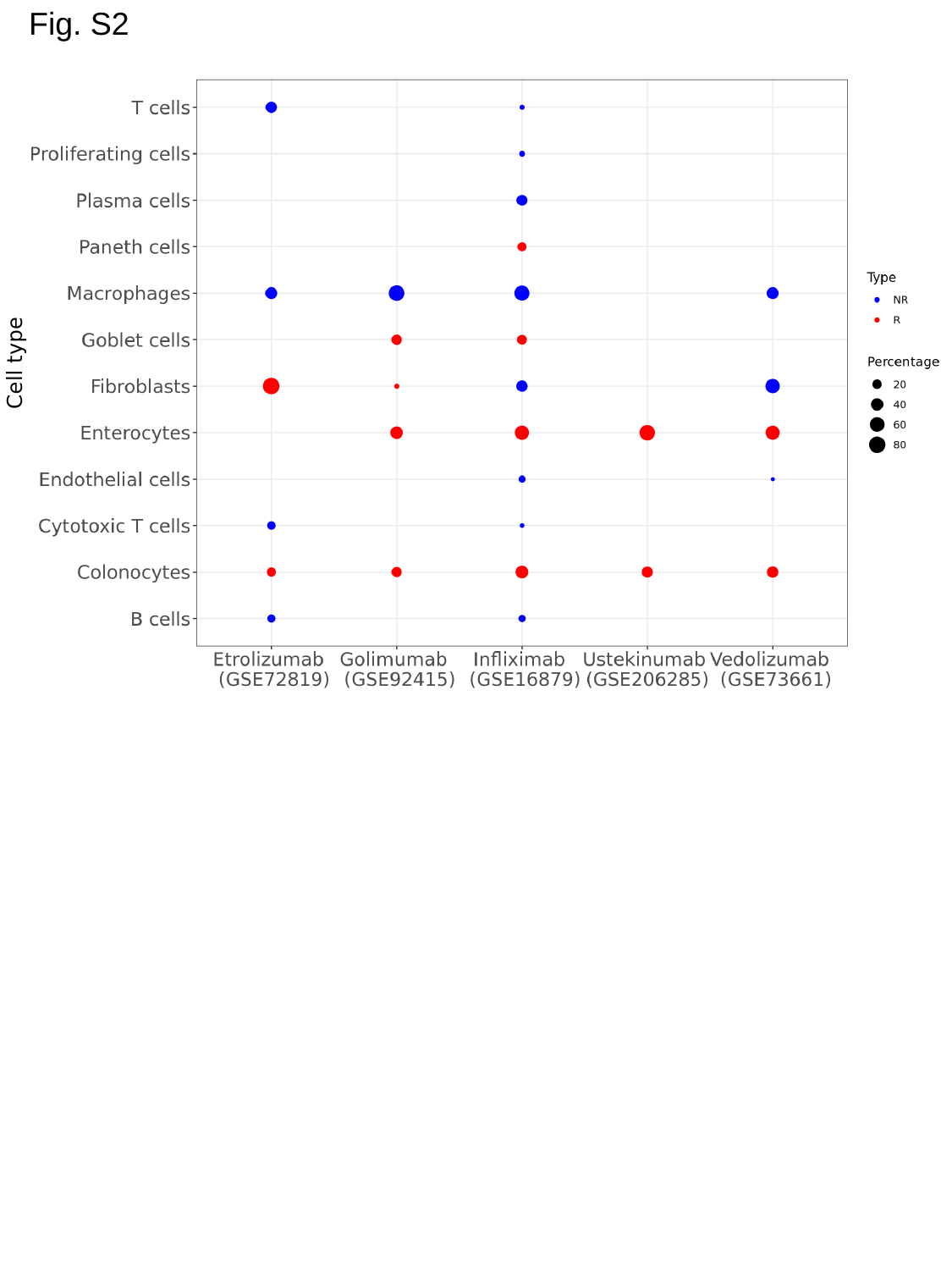

Fig. S2
